# Analysis of RNA sequence and structure in key genes of *Mycobacterium ulcerans* reveals conserved structural motifs and regions with apparent pressure to remain unstructured

**DOI:** 10.1101/2021.11.23.469657

**Authors:** Warren B. Rouse, Jessica Gart, Lauren Peysakhova, Walter N. Moss

## Abstract

Buruli Ulcer is a neglected tropical disease that results in disfiguring and potentially dangerous lesions in affected persons across a wide geographic area, which includes much of West Africa. The causative agent of Buruli Ulcer is *Mycobacterium ulcerans*, a relative of the bacterium that causes tuberculosis and leprosy. Few therapeutic options exist for the treatment of this disease beyond the main approach, surgical removal, which is frequently ineffective. In this study we analyze six genes in *Mycobacterium ulcerans* that have high potential of therapeutic targeting. We focus our analysis on a combined in silico and comparative sequence study of potential RNA secondary structure across these genes. The end result of this work was the comprehensive local RNA structural landscape across each of these significant genes. This revealed multiple sites of ordered and evolved RNA structure interspersed between sequences that either have no bias for structure or, indeed, appear to be ordered to be unstructured and (potentially) accessible. In addition to providing data that could be of interest to basic biology, our results provide guides for efforts aimed at targeting this pathogen at the RNA level. We explore this latter possibility through the *in silico* analysis of antisense oligonucleotides that could be used to target pathogen RNA.

**Author Summary:** Buruli Ulcer is a neglected tropical necrotizing skin disease endemic to West Africa and several other developing countries. The disease is known to be caused by *Mycobacterium ulcerans*, but the mode of transmission is not well understood. Here, we present findings on the RNA secondary structural landscape of key genes found in its genome and virulence plasmid. We also suggest potential therapeutic strategies to treat this disease that leverage a better understanding of RNA secondary structure. In our analysis we have predicted regions within these genes that are potentially ordered by evolution to have unusual structural stability and likely functionality, as well as regions that lack stable structure and may be unordered for accessibility. These structured regions can act as potential targets of both antisense oligonucleotide and small molecule therapeutics, while the unstructured regions may be most advantageous for only antisense oligonucleotides. Both strategies have been proven to be effective in other bacterial and viral pathogens; therefore, adaptation to this neglected disease may prove beneficial to developing more effective and efficient treatment options. Through our analysis of the RNA secondary structure landscape of key genes in *M. ulcerans*, we hope to provide other researchers with new avenues for development of novel therapeutic strategies to treat this devastating and neglected disease.

## Introduction

Buruli Ulcer (BU) is a skin-related neglected tropical disease (NTD) caused by the family of bacteria that also causes tuberculosis and leprosy (1). Scientifically called *Mycobacterium ulcerans* (*M. ulcerans*), this disease mainly affects the skin and, on rare occasions, bone. BU has been reported from 33 countries, 14 of which regularly report to the World Health Organization (2). BU is a disease that mainly affects West Africa and other tropical areas in the Western Pacific (3). *M. ulcerans* thrives in temperatures between 29–33 °C and requires a low oxygen concentration of 2.5% (4, 5), thus, the bacteria is able to easily spread in areas near the equator (3, 6). Limited knowledge on the mode of transmission of this pathogen makes it a critical target for further investigation (3).

BU relies on the virulence factor mycolactone for pathogenesis (7, 8). Mycolactone, an exotoxin secreted by *M. ulcerans* bacteria, has three major biological functions: cytotoxicity, immunosuppression, and analgesic effects (9, 10). These functions parallel the characteristics of the disease with large-scale skin ulceration, limited local inflammatory response, and limited or no pain associated with the damage, respectively (9, 11). *M. ulcerans* ability to block and degrade the nerves of infected tissue, and therefore inhibit the feeling of pain, makes it particularly dangerous (12). Due to the long incubation period and lack of symptoms, the infected often unknowingly carry the disease, prohibiting vital early diagnosis. As a result, most people affected by the bacteria do not know to seek treatment until it is too late (7).

Current treatment options include surgical removal of the skin, oral antibiotics, or dressing of the skin to help heal the wound (13). Surgery and skin grafting is only effective for early cases of BU because small areas of the skin are affected and can be safely removed (14). However, surgery is not a safe solution for cases that are diagnosed at a later stage of the disease because of the large area of the skin that is damaged (14). Rifampin, clarithromycin, and streptomycin are three antibiotic options that have been utilized to treat BU (14). However, oral antibiotics have not proven to be an ideal treatment, and patients who take these medications are frequently hospitalized because of adverse effects (7).

Very few studies have been conducted on finding novel treatments of BU. Since it only affects a small portion of the world (6, 14), therapeutics would have a relatively small return on investment—thus leaving BU a neglected disease. RNA-targeting therapeutics are emerging as exciting new therapies, which can take the form of small-molecules (15–17) or <u>a</u>nti<u>s</u>ense <u>o</u>ligonucleotides (ASOs; (18, 19)). The ASO approach could be particularly well suited as a strategy for treating BU or other neglected diseases, due to their rapid and, relatively, low cost to develop (20, 21). Additionally, as RNA-therapies grow in popularity, the projected future cost associated with their deployment is expected to decline (22).

ASOs can target RNA via sequence complementarity and prevent protein production by blocking the ribosome, masking functional elements, or degrading the transcript (23). Significant to this current study, ASOs have been previously used to target bacteria (19, 23), which offers hope that this method could be applied to BU. Targeting of RNA, both using small-molecules and ASOs, is greatly facilitated by analyses of target secondary and tertiary structure (17). Analysis of RNA structure has proven to be an important component of gene regulation in various human and pathogen genomes and genomes (24–27); however, to date, no analyses of RNA structure have been performed in *M. ulcerans*. A better understanding of RNA structure in this pathogen would thus provide information that could prove useful in treating BU, as it has been shown to be important to proper regulation of translation in other bacterial strains (28). In this current study we apply an in silico and sequence analysis pipeline for secondary structural RNA analysis, `ScanFold`, that is based on the identification of small local motifs with an unusually high propensity for ordered thermodynamic stability (29)—as measured by greater stability in evolved/ordered sequences vs in silico randomized sequences with the same nucleotide composition. This approach was optimized on viral human pathogens: HIV, Zika, human herpes viruses and, most recently, SARS-CoV-2 (24). Significantly, the latter analysis of SARS-CoV-2 led to the discovery (or highlighting) of structural motifs that are being targeted with both small molecules (30–32) and ASOs (33, 34). In this current analysis of *M. ulcerans*, we enhance our approach to make predictions that are of particular relevance to ASO identification.

## Methods

The *M. ulcerans* agy99 genome (NC_008611.1) and associated gff3 data as well as the pMUM001 virulence plasmid genome (NC_005916.1) and associated gff3 data were downloaded from NCBI (**S1 File**). Based on previous work done by Butt et al. (35), six essential genes with no host homology were chosen for analysis. The chosen target genes are *Mul_RS01615*, *Mul_RS01365*, *Mul_RS04730*, *Mul_RS09540*, *Mul_RS04200* and *Mul_RS00210*, or Targets 1-6 respectively. All genes except *Mul_RS00210* were found in the NC_008611.1 genome and *Mul_RS00210* was found in the NC_005916.1 genome.

We loaded these genomes and associated gff3 files into `IGV-ScanFold` (version 0.2.0) (https://github.com/ResearchIT/IGV-ScanFold/releases/) to view the RNA sequence of *M. ulcerans* as a whole. After searching for the genes, they were initially scanned in their entirety by highlighting the gene and using the “ScanFold>Run ScanFold on selected region” option. For this analysis the default parameters were changed to a 120 nt window size, a 1 nt step size, 100 randomizations per window, mononucleotide shuffling, 37 °C temperature, competition of 1 (to demand that only one unique base pair per nucleotide is possible), either forward or reverse strand depending on the gene polarity, and `RNAfold` folding algorithm were selected. Metrics obtained from `ScanFold` include the MFE (a measure of thermodynamic stability), ΔG z-score (a measure of ordered stability that can indicate potential function), and ensemble diversity or ED (a measure of predicted structure’s conformational volatility). The MFE is estimated by the predicted Gibb’s folding free energy change (the ΔG°) going from a fully denatured (random coil) RNA to an ordered 2D structure, where more negative values indicate increasing favorability and greater stability. ΔG z-scores identify structures that have propensity for ordered stability, where negative values indicate the number of standard deviations more stable the native sequence is compared to any randomized version of it (36, 37). ED uses the RNA partition function to compare the distance between the optimal thermodynamic structure and suboptimal conformations (38). This distance is measured as the number of base pairs different between Boltzmann weighted conformations and is averaged across the ensemble. Lower ED values indicate a single dominant conformation, while higher EDs suggest multiple conformations or a lack of defined structure (39). Additionally, arc diagrams depicting weighted z-score structures are produced where blue, green, and yellow arcs indicate z-scores ≤ −2, ≤ −1, and <0, respectively. For more information on these metrics, see the ScanFold methods paper (40). After initial analysis, several genes were found to have regions of low z-score near the 3’ and 5’ ends. All genes were re-scanned with an additional 50-300 nucleotides upstream and downstream of the start and stop codon, respectively. This allowed us to note any patterns in stability immediately outside the targets.

To further assess the potential functionality of structures found in these genes, covariation analysis was completed using the CM-Builder analysis pipeline (41). All structures with a z-score of −2 or less were analyzed for covariation using the `cm-builder` perl script (41). This script builds off the `RNAFramework` toolkit (42, 43) and utilizes Infernal (here using release 1.1.2; (44)) to build and search for covariation models from each predicted ScanFold secondary structure. Briefly, `cm-builder` uses `Infernal` to generate a covariance model (CM) using the sequence and structure of the selected motif. A database of sequences is built for `Infernal` using a BLAST search (45). For *M. ulcerans*, the nt collection database was searched for each of the selected motifs found in the target genes using following parameters: BLASTn (Somewhat similar), Organism (bacteria, taxid:2), and a max sequence number of 5000. `Infernal` then takes the BLAST data and aligns the sequences removing any redundant and severely truncated alignments. The resulting set of homologs from the alignment of BLAST hits is then aligned to the original CM, which is used to build a new CM. The whole process is repeated three times. Finally, the resulting alignment is refactored to remove gap-only positions and include only bases spanning the first to the last base-paired residue. The final alignment file is analyzed using `R-scape` (42) where APC corrected *G*-test (46) statistics are used to identify motifs showing significantly covarying base-pairs using the default E value of 0.05. Statistically significant covariation is indicative of evidence for a conserved RNA base pair. For *M. ulcerans*, these statistically significant base pairs were then annotated on the 2D models made using `VARNA` (47).

After running `ScanFold` to define regions of unusual thermodynamic stability, and `cm-builder` to identify statistically significant covarying base pairs, the sequences of each gene were loaded into `OligoWalk` (48) and ran with default parameters (Mode: Break Local Structure, Oligo Chemistry: DNA, Oligo Length: 18nt, and Oligo Concentration: 1uM). The overall ΔG, duplex ΔG, intra-oligo ΔG, and inter-oligo ΔG values for 18 nucleotide complementary oligonucleotides across each gene sequence was obtained for further analysis. Additionally, using the sequence fasta file, final partners file, and log files from `IGV-ScanFold` an in-house python script (https://github.com/moss-lab/ScanFold_Oligo_Metrics/blob/main/ScanFold_Oligo_Metrics.py) was used to partition the data into 18nt fragments and determine the average ED, MFE, and z-score for each. This was accomplished using a tiled window approach, which allowed for the complementary analysis of `ScanFold` and `OligoWalk` metrics for the same fragments. For more information on the data generated by this script see the ReadMe file on GitHub. Using `Oligowalk` and `ScanFold` data, bar graphs were made for every gene (**S2 File**). All data used to create these graphs and analyze the results, can be found in (**S3 File**). Following the work of Matveeva et. al., oligonucleotides that had intra-oligo values greater than −8 kcal/mol, inter-oligo values greater than −1.1 kcal/mol, duplex ΔG values of less than −15 kcal/mol, and negative overall ΔG values were considered as optimal ASO targets to begin further analysis. This data was then compared to the ScanFold fragment data and trends were noted.

## Results

We focused on six genes found to be essential to *M. ulcerans* (35, 49) with no known homology to human hosts: *Mul_RS01615*, *Mul_RS01365*, *Mul_RS04730*, *Mul_RS09540*, *Mul_RS04200* and *Mul_RS00210*. These were selected for their essentiality to the *M. ulcerans* and their homology to genes targeted in other pathogens (35, 49). These were subjected to the full `ScanFold` pipeline as integrated in `IGV-ScanFold`. In the first step `ScanFold-Scan` was used to define the local RNA folding landscape of each gene (**S4 File**). Here, a scanning analysis window of 120 nucleotides (nt) was moved one base at a time across each gene sequence while predicting several folding metrics: the minimum free energy of folding (MFE, ΔG°), its associated base pairing model (secondary structure), and a thermodynamic z-score that compares the MFE of the natively ordered RNA to randomized version to identify propensity for ordered stability as indicated by negative values; details on all metrics in the Methods Section. In the second stage, `ScanFold-Fold`, folding metrics are partitioned to each nucleotide and base pair, and consensus base pairs across all windows— weighted by their propensity for ordered stability (thermodynamic z-score) —are identified and output as distinct motifs (**S5 File**).

A summary of results across each gene is seen in **Table 1**. Here, we see that average MFE ΔG values across all analyzed genes ranged from −48.40 kcal/mol to −33.23 kcal/mol. This difference in predicted RNA folding stability is directly correlated with GC%, as expected, which ranged from 68.14%to 61.32%. The average z-score and ensemble diversity (ED), however, do not follow trends for ΔG or GC%. The z-score quantifies how much greater-than-random the folding stability of an RNA sequence is, which is primarily dependent on the sequence order and not its nucleotide composition. Likewise, the ED value indicates the diversity of potential structural conformations in an RNA’s folding ensemble, which also appears to be an evolved property of ordered/functional RNA sequences (39). The average z-score ranged from −0.82 to −0.03, while the average ED ranged from 20.64 to 26.85. Here we see that the gene with the lowest (most favorable) ΔG and highest GC% (*Mul_RS01615*) has the highest (least favorable) average z-score and highest ED. The gene with the lowest average z-score (*Mul_RS04730*) also showed the highest percent of windows with z-score <−1 (39.55%). This did not hold true for z-scores <−2 (two standard deviations more stable than random), but the gene with the second lowest average z-score (*Mul_RS01365*) showed the highest percent of windows with z-scores <−2 (14.76%). The gene with the smallest fraction of its nucleotides spanned by low z-score windows was *Mul_RS01615* (19.0% and 4.4% for the −1 and −2 z-score cutoffs, respectively).

**Table 1.**
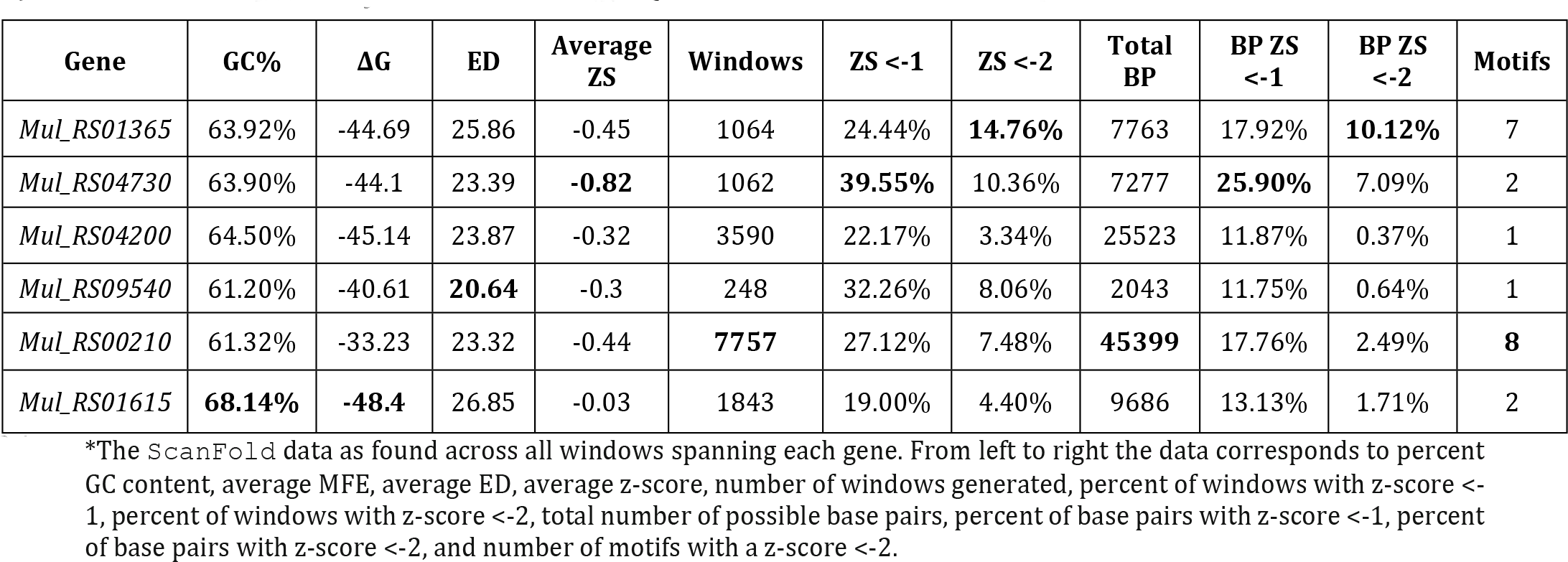
Summary of ScanFold data across each target gene.*

| Gene | GC% | $\Delta G$ | ED | Average ZS | Windows | ZS <-1 | ZS <-2 | Total BP | BP ZS <-1 | BP ZS <-2 | Motifs |
| --- | --- | --- | --- | --- | --- | --- | --- | --- | --- | --- | --- |
| *Mul_RS01365* | 63.92% | -44.69 | 25.86 | -0.45 | 1064 | 24.44% | **14.76%** | 7763 | 17.92% | **10.12%** | 7 |
| *Mul_RS04730* | 63.90% | -44.1 | 23.39 | **-0.82** | 1062 | **39.55%** | 10.36% | 7277 | **25.90%** | 7.09% | 2 |
| *Mul_RS04200* | 64.50% | -45.14 | 23.87 | -0.32 | 3590 | 22.17% | 3.34% | 25523 | 11.87% | 0.37% | 1 |
| *Mul_RS09540* | 61.20% | -40.61 | **20.64** | -0.3 | 248 | 32.26% | 8.06% | 2043 | 11.75% | 0.64% | 1 |
| *Mul_RS00210* | 61.32% | -33.23 | 23.32 | -0.44 | **7757** | 27.12% | 7.48% | **45399** | 17.76% | 2.49% | **8** |
| *Mul_RS01615* | **68.14%** | **-48.4** | 26.85 | -0.03 | 1843 | 19.00% | 4.40% | 9686 | 13.13% | 1.71% | 2 |

\*The ScanFold data as found across all windows spanning each gene. From left to right the data corresponds to percent GC content, average MFE, average ED, average z-score, number of windows generated, percent of windows with z-score <-1, percent of windows with z-score <-2, total number of possible base pairs, percent of base pairs with z-score <-1, percent of base pairs with z-score <-2, and number of motifs with a z-score <-2.

In the prediction of the MFE ΔG for each analysis window, a model secondary structure is also generated. Across each gene this resulted in many predicted base pairs, where a specific nucleotide can be paired differently across several overlapping windows. As a result, many potential base pairing partners may be predicted per nucleotide. `ScanFold-Fold` predicts a single structural context (paired or unpaired) for each nucleotide based on its contributions to low z-score windows (indicating ordered stability and, potentially, function). The genes that had the greatest percentages of low z-score base pairs were *Mul_RS04730* and *Mul_RS01365*; *Mul_RS04730* had 25.90% of its base pairs predicted with average z-score <−1 and *Mul_RS01365* had 10.12% of its base pairs predicted with average z-score <−2 (**Table 1**). Additionally, `ScanFold-Fold` extracts discrete structural motifs (i.e., single hairpins or multi-branched stem loops) containing at least one base pair with an average z-score <−2. This results in a list of motifs for each gene, where the longest gene *Mul_RS00210* had the greatest number of motifs (8 motifs) and the much shorter gene *Mul_RS01365* had the second greatest number of motifs (7 motifs).

In summary, all predictions indicate a particular importance for functional RNA secondary structures encoded within the genes for virulence factor production and cell wall biosynthesis. The `ScanFold-Fold` results also give us a means of generating motifs of interest for further analysis.

### Results from analysis of RNA folding in *Mul_RS01365*

*Mul_RS01365* encodes a protein that is homologous to the *desA2* (stearoyl-ACP desaturase) protein of *M. tuberculosis*. It is thought to function via the catalysis of the initial conversion of saturated fatty acids to unsaturated fatty acids in lipid metabolism.

A summary of `ScanFold` data is shown on **Fig. 1**, where blue, green, and yellow arcs indicate base pairing with average z-scores ≤ −2, ≤ −1, and <0, respectively. Interesting trends are seen in the `ScanFold-Scan` folding metrics partitioned per nucleotide of the transcript. The overall thermodynamic stability remains flat across *Mul_RS01365* until scans reach the 3′ end of this RNA, where stability increases (more negative MFE ΔG° in windows overlapping these nucleotides). Despite fairly monotonous trends in ΔG°, evidence for regions of ordered RNA stability (negative z-scores) are clustered into two distinct regions at the 5′ and 3′ ends of the coding region, respectively. Interestingly, the core coding region is spanned by a region with positive z-scores, which indicates that the ordered/evolved sequence is less stable than predicted based on nucleotide content: i.e., the evolved sequence may be ordered to be unstructured or accessible. This region of unusual instability correlates with spikes in the ensemble diversity, which indicates a volatile ensemble of potential RNA secondary structures or a lack of stable structure. Conversely, the low z-score clusters overlap regions of low ensemble diversity, indicating one (or a few similar) conformation(s).

**Fig 1.**
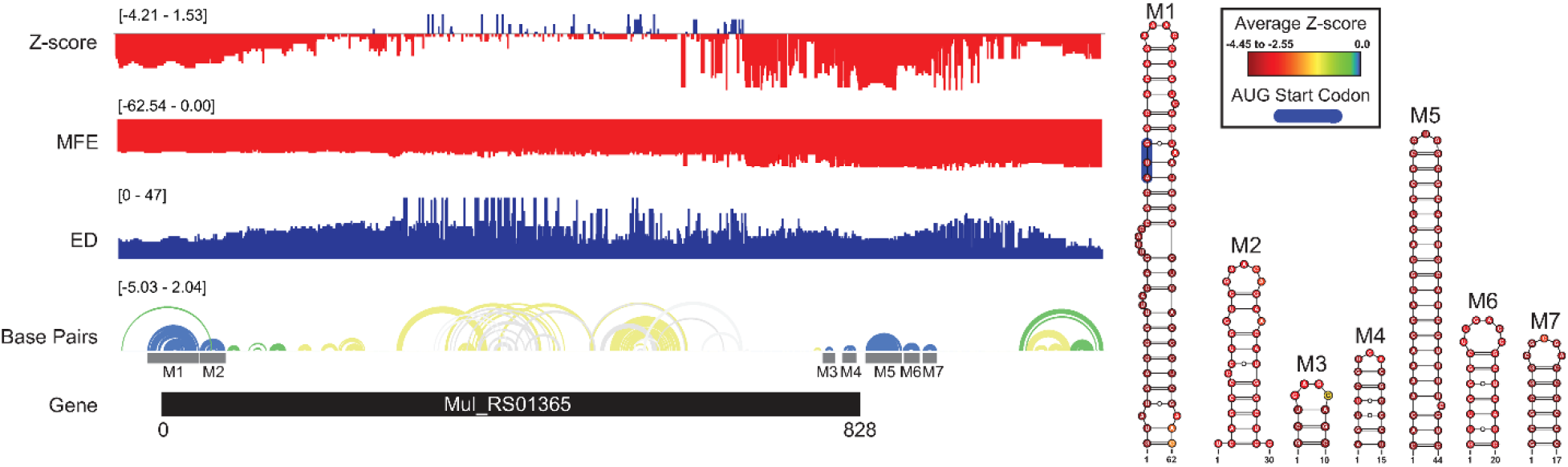
*Mul_RS01365* ScanFold results and 2D models. Global `ScanFold` results for *Mul_RS01365*. The ∆G z-score, minimum free energy (MFE), ensemble diversity (ED), gene cartoon, and base pair arc diagram (top to bottom) are shown to the left. All −2 ∆G z-score structures found are represented as 2D models to the right. The base pair arc diagram is annotated with gray boxes to show the location of M1-7 across the gene. In the 2D structure model of M1, the canonical AUG start codon is annotated in blue. The nucleotides of each structure are annotated with the average per nucleotide z-scores where the most negative are indicated in red and the most positive are indicated in blue.

When the low z-score windows were evaluated by `ScanFold-Fold`, seven distinct <u>m</u>otifs (M1-7) were identified with exceptionally low (<−2) z-score-weighted base pairs (**Fig. 1**). These motifs comprise two upstream hairpins (M1 and M2) and five downstream hairpins that span the start and stop sites of translation, respectively. M1 is notable for containing the start codon embedded within a stable helix and for being the longest thermodynamically stable hairpin predicted for this gene. When evaluated for their conservation across mycobacterial species, none of the proposed base pairs were found to have statistically significant covariation (**S6 File**). The proposed structures were, however, found to be present/conserved across a wide array of species.

In M1, for example, the hairpin structure is preserved across pathogenic species of mycobacteria (**Fig. 2**). Conservation is highest in the terminal hairpin loop region that contains the start codon, and trails off toward the basal stem, where deletions and inconsistent mutations (that ablate canonical base pairing potential) would be predicted to weaken the basal stem. The core hairpin is best preserved (100% preservation of base pairing) in the medically significant species, *M. gordonae*, *kansasii*, *tuberculosis* and *bovis*, while inconsistent mutations begin accumulating in *M. fortuitum,* where a U>C mutation disrupts a single AU pair (**Fig. 2**). M. avium had the most inconsistent mutations in the core hairpin, that would be predicted to weaken or break four base pairs (out of 15 base pairs). Conversely, while no compensatory mutations are observed in the core hairpin, six species had at least one consistent change, where single point mutations preserve base pairing; *M. Leprae* had two consistent mutations, however, these were offset by several inconsistent changes.

**Fig 2.**
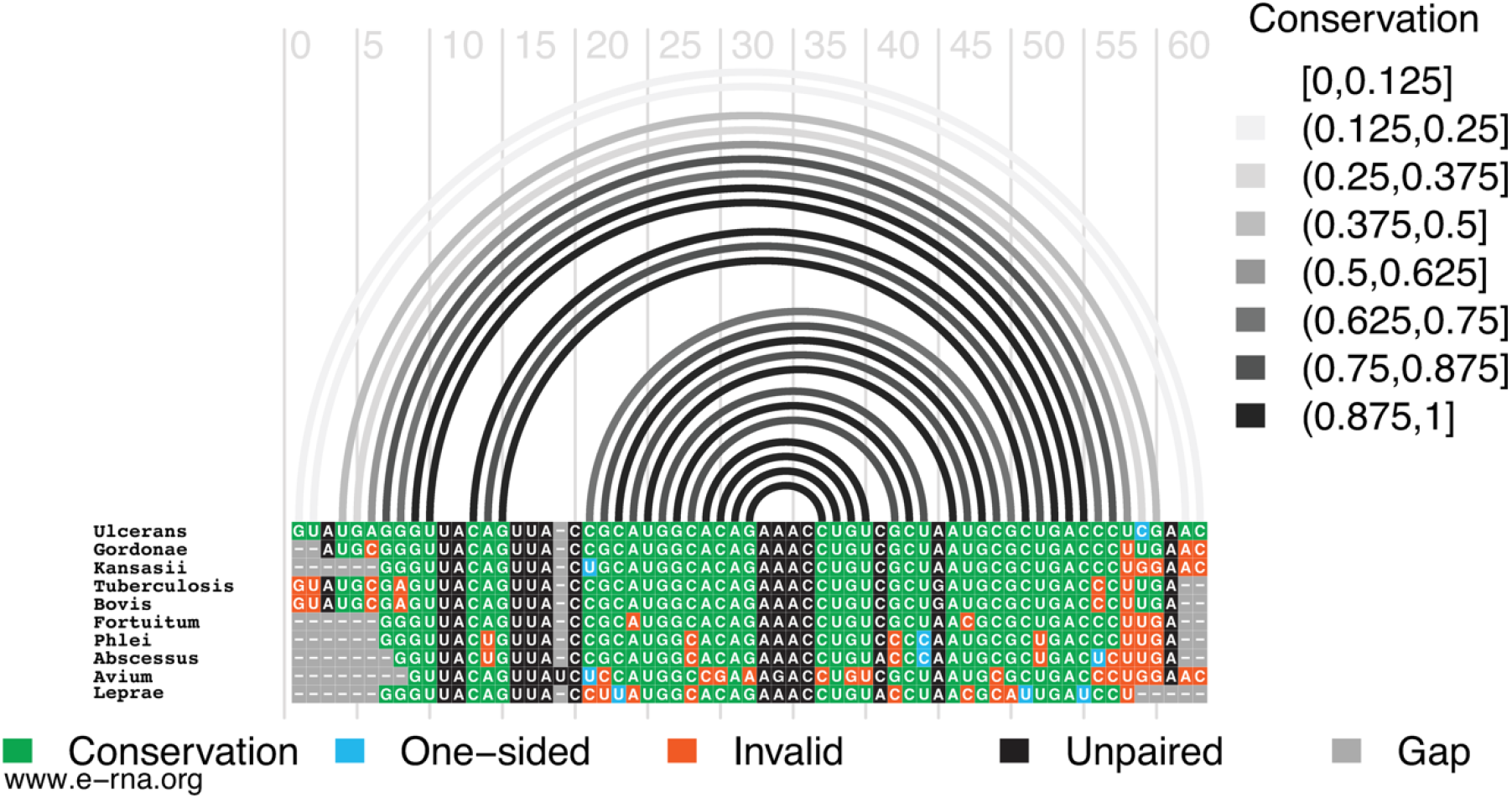
Conservation of *Mul_RS01365* Motif 1. Conservation of sequence and structure across pathogenic mycobacterium strains M. ulcerans*, gordonae*, *kansasii*, *tuberculosis*, *bovis*, *fortuitum, phlei, abscessus, avium, and leprae*. Boxes are color coded based on their representation. Conservation (green), covariation (dark blue), consistent mutations (light blue), inconsistent mutations (orange), unpaired nucleotides (black), and alignment gap (grey).

### Results from analysis of RNA folding in *Mul_RS04730*

*Mul_RS04730* encodes a protein homologous to the *rpoA* (alpha chain of RNA polymerase) of *M. tuberculosis*. This protein functions as a component of the DNA-dependent RNA polymerase responsible for bacterial genome replication.

A summary of `ScanFold` data is shown in **Fig. 3** following the same color scheme mentioned previously. Trends are seen in the `ScanFold-Scan` folding metrics partitioned per nucleotide of the transcript. The overall thermodynamic stability (ΔG°) remains flat across the entirety of the transcript. Despite the flatline trend in ΔG°, evidence for regions of ordered RNA stability (negative z-scores) are clustered into one distinct region just upstream of and at the 5′ end of the coding region. The coding region is predominantly spanned by structures with z-scores of 0 to −1 with a few structures of ≤ −2 and ≤ −1 interspersed. This indicates that the ordered/evolved sequence is overall only slightly more stable than predicted based on nucleotide content: i.e., the evolved sequence may be ordered to be structured. The regions of only slightly increased stability correlate with spikes in the ensemble diversity when compared to those of the much more stable region just upstream of and at the 5′ end of the coding region. This indicates a volatile ensemble of potential RNA secondary structures or a lack of stable structure across the core coding sequence; whereas the more 5’ region shows a less volatile ensemble or more stable structures that may form potentially functional structures.

**Fig 3.**
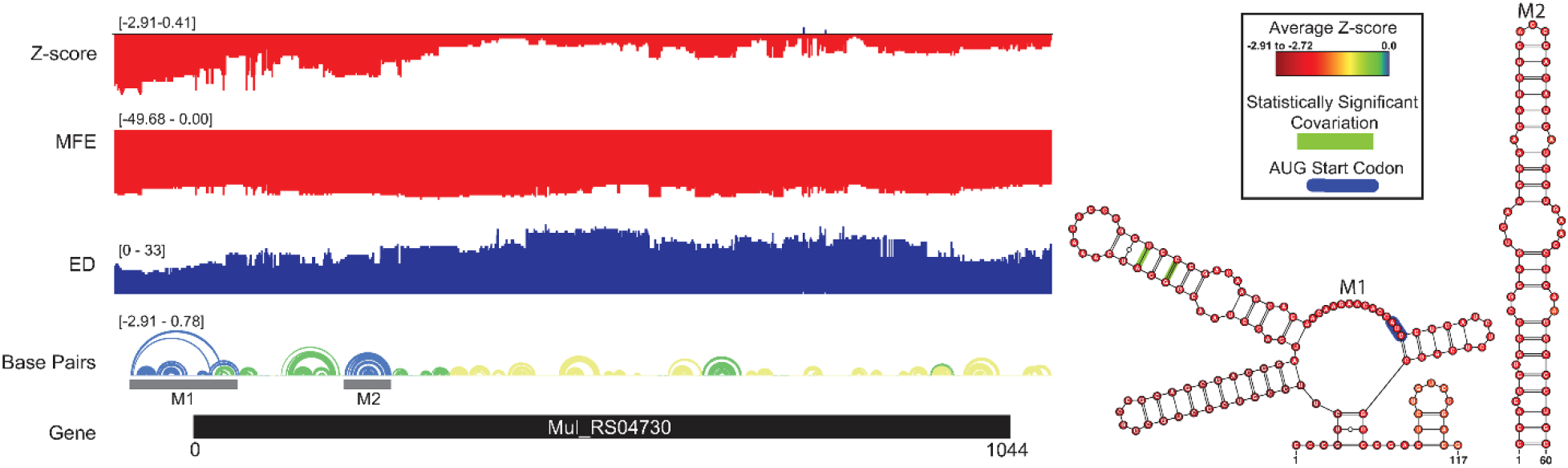
*Mul_RS04730* ScanFold results and 2D models. Global `ScanFold` results for Mul_RS04730. The ∆G z-score, minimum free energy (MFE), ensemble diversity (ED), gene cartoon, and base pair arc diagram (top to bottom) are shown to the left. All −2 ∆G z-score structures found are represented as 2D models to the right. The base pair arc diagram is annotated with gray boxes to show the location of M1-2 across the gene. In the 2D structure model of M1, the canonical AUG start codon is annotated in blue. Nucleotides that exhibit statistically significant covariation are annotated by green bars across the base pair. The nucleotides of each structure are annotated with the average per nucleotide z-scores where the most negative are indicated in red and the most positive are indicated in blue.

When the low z-score windows were evaluated by `ScanFold-Fold`, two distinct <u>m</u>otifs (M1-2) were identified with exceptionally low (<−2) z-score-weighted base pairs (**Fig. 3**). These motifs comprise one upstream multi-branch helix (M1) and one downstream hairpin (M2) that spans the start site of translation and is a part of the coding sequence, respectively. M1 is notable for containing the start codon in a single stranded region between two hairpins of the multi-branch helix. When evaluated for their conservation across mycobacterial species, two of the proposed base pairs were found to have statistically significant covariation (**S6 File** and **Fig. 3**). In addition, part of the M1 structure and all of the M2 structure were found to be present/conserved across a wide array of species.

In M1, for example, the hairpin structure is preserved across pathogenic species of mycobacteria (**Fig. 4**). Conservation is highest in the central hairpin that shows evidence of significant covariation and the downstream hairpin containing the start codon. However, toward the 5’ end of the structure, conservation drops off, as homologous sequences were not found here. The small hairpin containing the last nucleotide of the start codon is best conserved (100% preservation of base pairing) in the medically significant species, *M. gordonae*, *kansasii*, *fortuitum, phlei, abscessus, and avium*; while inconsistent mutations begin accumulating in M. *tuberculosis, bovis, and leprae,* where a U>A mutation disrupts a single AU pair (**Fig. 4**). *M. gordonae* had the most inconsistent (with proposed 2D structure) nucleotides in the 5’ hairpin (10 mutations), while *M. tuberculosis* and *bovis* had the most inconsistent nucleotides in the core hairpin (5 mutations). Interestingly, two compensatory mutations are observed in the core hairpin, which match those found in the `cm-builder` analysis. Seven of the ten species had at least one consistent change, where single point mutations preserve base pairing; *M. kansasii* had two consistent mutations, however, these were offset by several inconsistent changes. Notably, *M. leprae*, another necrotizing bacterium, appeared to have the highest level of conservation across the entire structured region when compared to *M. ulcerans*. Overall, the core hairpin and next downstream hairpin containing the start codon appear to be quite conserved, and when analyzed against all other data seems to indicate this region could be potentially functional.

**Fig 4.**
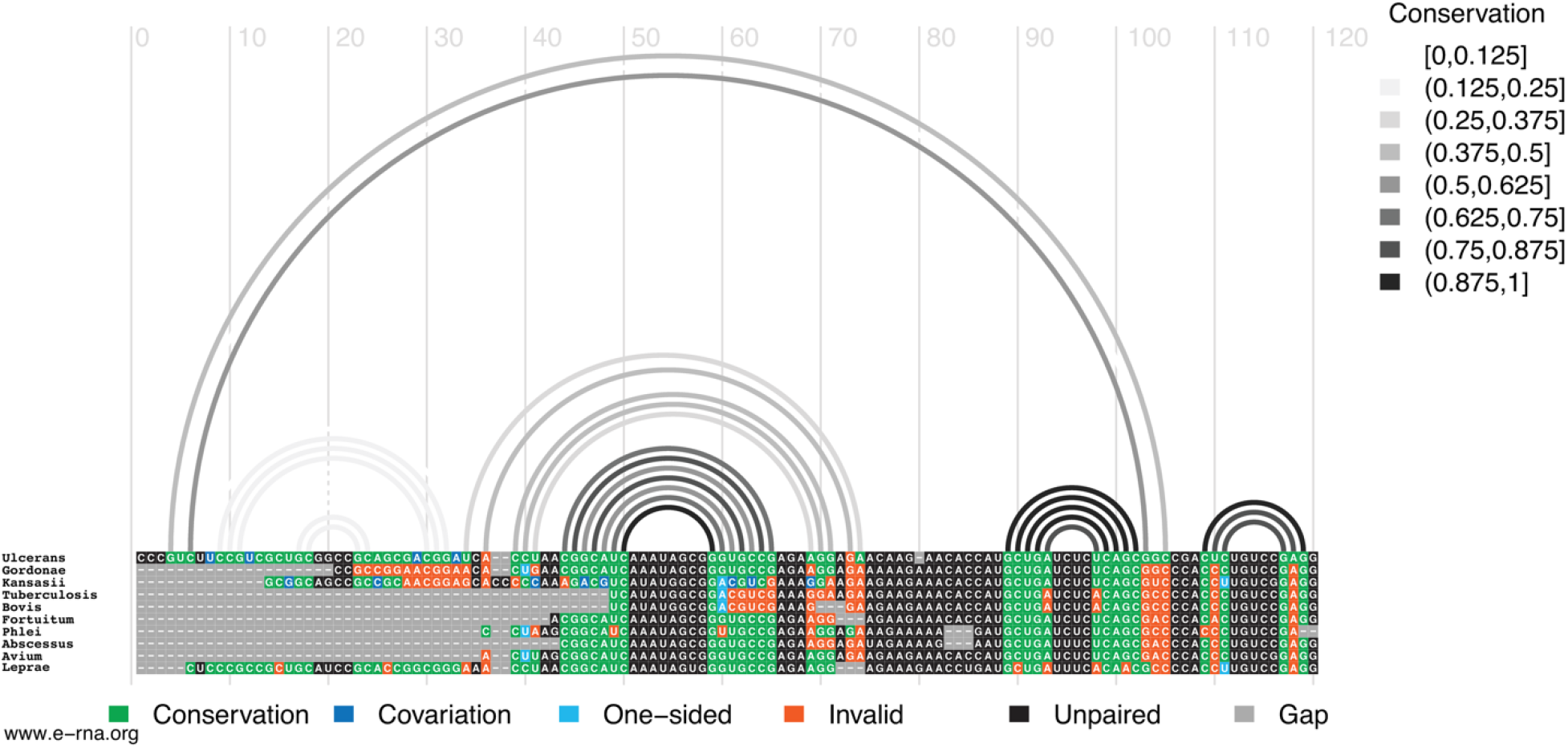
Conservation of *Mul_RS04730* Motif 1. Conservation of sequence and structure across pathogenic mycobacterium strains M. ulcerans*, gordonae*, *kansasii*, *tuberculosis*, *bovis*, *fortuitum, phlei, abscessus, avium, and leprae*. Boxes are color coded based on their representation. Conservation (green), covariation (dark blue), consistent mutations (light blue), inconsistent mutations (orange), unpaired nucleotides (black), and alignment gap (grey).

### Results from analysis of RNA folding in *Mul_RS04200*

*Mul_RS04200* encodes a protein homologous to the *rpoB* (beta chain of RNA polymerase) of *M. tuberculosis*. This protein functions as a component of the DNA-dependent RNA polymerase responsible for bacterial genome replication.

Interesting trends are seen in the `ScanFold-Scan` folding metrics partitioned per nucleotide of the transcript (**S1 Fig.**). The overall thermodynamic stability (ΔG°) remains flat across the entirety of the *Mul_RS04200* gene. Despite the flatline trend in ΔG°, there is evidence for regions of ordered RNA stability (negative z-scores). Z-scores remained relatively negative across the entire gene, but regions of lower z-scores were noted. One region in the middle of the gene did show a significant decrease in z-score below −2, while the majority of the gene’s 3’ end displayed lower than average z-scores (indicating increased stability towards the end of the gene). In contrast to the 3’ end, a region of positive z-scores is observed approximately 750 nucleotides from the 5’ end.

The `ScanFold-Fold` motif found in the lowest z-score region (M1; **S1 Fig.**) had base pairs with significantly low z-scores (< −2) which increased upstream and downstream of the hairpin. Notably, as base pairing extended out from the basal stem, the z-scores steadily increased until the final two bulges and terminal loop became only slightly negative.

### Results from analysis of RNA folding in *Mul_RS09540*

*Mul_RS09540* encodes a protein homologous to the *rpoZ* (omega component of RNA polymerase) of *M. tuberculosis*. This protein functions as a component of the DNA-dependent RNA polymerase responsible for bacterial genome replication.

Interesting trends are seen in the `ScanFold-Scan` folding metrics partitioned per nucleotide of the gene (**S2 Fig.**). Compared to the others, *Mul_RS09540* is the smallest gene that was analyzed. The overall thermodynamic stability (ΔG°) remains uniform across the entirety of the gene. *Mul_RS09540* also had the fewest base pairs with < −2 z-score. Near the 3’ end, there are two distinct clusters of z-score values that increase into the positive range, thus indicating a region that may be evolved to have reduced structural propensity. This same unstable region correlates with high ensemble diversity, further indicating the lack of ordered and stable structure at the 3’ end. Conversely, low z-score regions overlapped with regions of low ensemble diversity, indicating that stronger RNA secondary structures are present near the core of the coding sequence. Interestingly, the most stable area of the structure seems to occur towards the middle of the gene rather than near the 3’ or 5’ ends.

The `ScanFold-Fold` motif found for this region (M1; **S2 Fig**) is predicted to form a 106 nucleotide hairpin that displays interesting trends in stability. In this structure the most significantly low z-score base pairs were found in the first 5 base pairs of the basal stem. As base pairing extended out from the basal stem, the z-scores steadily increased until the final two bulges and terminal loop became only slightly negative.

### Results from analysis of RNA folding in *Mul_RS00210*

*Mul_RS00210* encodes a protein homologous to the *Pks7* (putative polyketide synthase) of *M. tuberculosis*. This protein is likely to function in lipid metabolism where it is believed to be involved in intermediate steps for the synthesis of a polyketide molecule, mycolactone. This polyketide has been shown to act as the virulence factor essential for infection and the painless nature of the ulcers (49). Rather than falling within the bacterial genome, the *Mul_RS00210* gene occurs within the 174 kb virulence plasmid pMUM001 that encodes a cluster of giant polyketide synthases (49). Until recently this was an uncharacterized example of plasmid-mediated virulence in a Mycobacterium, and it is believed that the pathogenicity of *M. ulcerans* is due the acquisition of pMUM001 by horizontal transfer (49).

A summary of `ScanFold` data is shown on (**Fig. 5**). The overall thermodynamic stability (ΔG°) of the *Mul_RS00210* transcript is much more variable than that of all other genes analyzed in this study. Evidence for regions of ordered RNA stability (negative z-scores) are clustered into eight distinct regions throughout the coding region. Here, there are eight structured regions with z-scores <−2 and five small regions of positive z-scores interspersed (the first region is in the intergenic region). This indicates that the ordered/evolved sequences making up these eight structures are much more stable than predicted based on nucleotide content: i.e., the evolved sequence may be ordered to have structure. These regions of high ordered stability (low z-score) correlate with dips in the ensemble diversity when compared to those found in regions of much lower ordered stability indicating that they are more stable and have a less volatile ensemble of potential RNA secondary structures that can form. The few regions of positive z-score may be indicative of regions that are evolved to be unstructured and therefore accessible to regulatory molecules.

**Fig 5.**
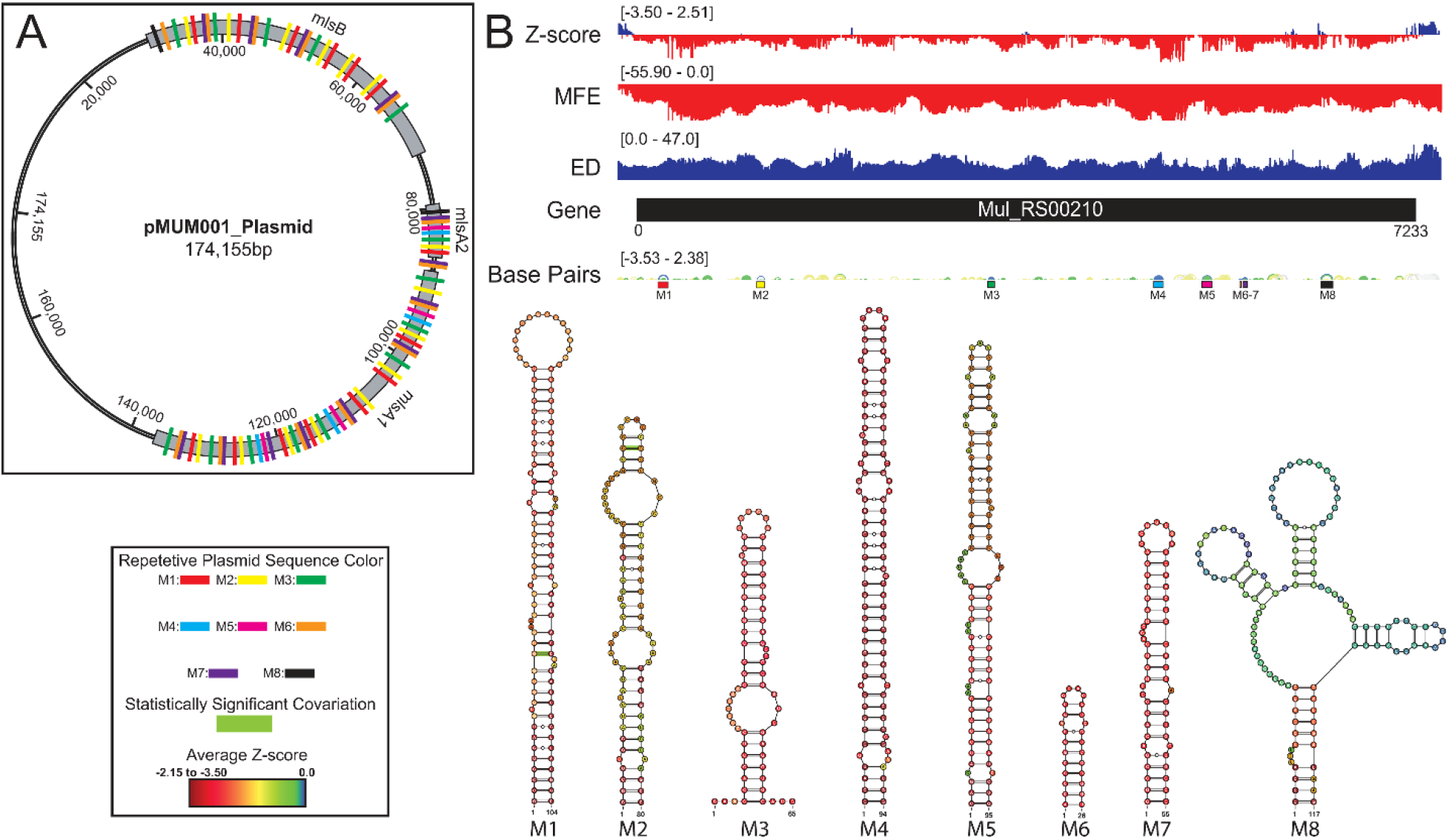
*Mul_RS00210* ScanFold results, 2D models, and structure occurrence throughout the pMUM001 plasmid. Global `ScanFold` results for *Mul_RS00210* including the location of repetitive sequence and structure elements across the pMUM001 virulence plasmid. A) A schematic diagram of the 174kb pMUM001 virulence plasmid with the locations of the three largest genes that encode the subunits responsible for production of mycolactone. The colored boxes annotated across these three genes represent the locations of the sequences that form structures M1 (red), M2 (yellow), M3 (green), M4 (blue), M5 (pink), M6 (orange), M7 (purple), and M8 (black) in the Mul_RS00210 gene. B) Global ScanFold results for Mul_RS00210 including (top to bottom) the ∆G z-score, minimum free energy (MFE), ensemble diversity (ED), gene cartoon, base pair arc diagram, and 2D models of −2 ∆G z-score structures. The base pair arc diagram is annotated with colored boxes (correlated to the colors in panel A) to show the location of M1-8 across the gene. In the 2D structure models, nucleotides that exhibit statistically significant covariation are annotated by green bars across the base pair. The nucleotides of each structure are annotated with the average per nucleotide z-scores where the most negative are indicated in red and the most positive are indicated in blue.

When the low z-score windows were evaluated by `ScanFold-Fold`, eight distinct <u>m</u>otifs (M1-8) were identified with exceptionally low (<−2) z-score-weighted base pairs (**Fig. 5**). These motifs comprise seven hairpins (M1-7) and one multi-branch helix (M8). Here, all the hairpins were found to have base pairs with significantly lower than average z-scores (<−2), whereas the multi-branch helix was found to only have significantly low z-scores in the basal stem. When evaluated for conservation across mycobacterial species, two of the proposed base pairs, found in structures (M1-2), were found to have statistically significant covariation (**Fig. 5** and **S6 File**). In addition, all structures were found to be present/conserved across a wide array of species

One unique feature of all −2 z-score structures found in the gene was their presence throughout all three genes encoding the polyketide synthase subunits responsible for production of mycolactone (**Fig. 5**). Predicted motifs are thus, replicated multiple times throughout the plasmid—up to 15 times (**Fig. 5**). Although this observed multiplication of structural elements is unique to our analysis, this finding is not entirely surprising as Stinear et. al found that these three genes have stretches of up to 27kb of near identical nucleotide sequence (99.7%). Additionally, of the 105-kb mycolactone locus, there is only ~9.5 kb of unique, nonrepetitive sequence (49).

### Results from analysis of RNA folding in *Mul_RS01615*

*Mul_RS01615* encodes a protein that is homologous to the *accD3* (putative acetyl CoA carboxylase carboxyl transferase-beta subunit) of *M. tuberculosis*. This protein is a component of the acetyl coenzyme A carboxylase complex and plays a functional role in lipid metabolism.

MFE folding stability varied across the gene, with a general trend toward lower stability (more positive values) toward the 3′ end (**Fig. 6**). A distinct cluster of low z-score nucleotides occurs in the core coding region of the gene, which overlaps a cluster of distinctly low ensemble diversity values. The `ScanFold-Fold` motif built for this region, M1, contains a multibranch loop structure formed of two hairpin loops with low, but not significantly negative, z-score nucleotides and base pairs. The multibranch loop structure sits atop a long stem formed by significantly low (<−1) z-score base pairs and nucleotides, where the basal stem (composed of six base pairs) had the lowest (<−2) z-scores. Though the ensemble diversity was low across M1, the region spanning the two hairpins of the multibranch loop were somewhat higher (**Fig. 6**), suggesting potential conformational dynamics for the two hairpins. Notably, the region immediately downstream of M1 was spanned by positive z-scores, indicating a potential for ordered *instability* of secondary structure. A second structural motif, M2, was predicted immediately downstream of the annotated open reading frame (ORF) for *Mul_RS01615* (**Fig. 6**). This motif falls in a somewhat diffuse region of low z-scores that overlaps a cluster of moderate ensemble diversity; thus, while the base pairs and nucleotides comprising M2 have significantly low z-scores (<−2), the predicted conformational ensemble does not appear to be particularly tight (i.e., potential for dynamics). It is worth noting that the transcript annotations for *M. ulcerans* are not sufficient to determine if this motif falls within the 3′ UTR *of Mul_RS01615*. While both motifs in *Mul_RS01615* were found to have potentially homologous sequences and structures in other mycobacteria (**S6 File**), significant covariation was not observed.

**Fig 6.**
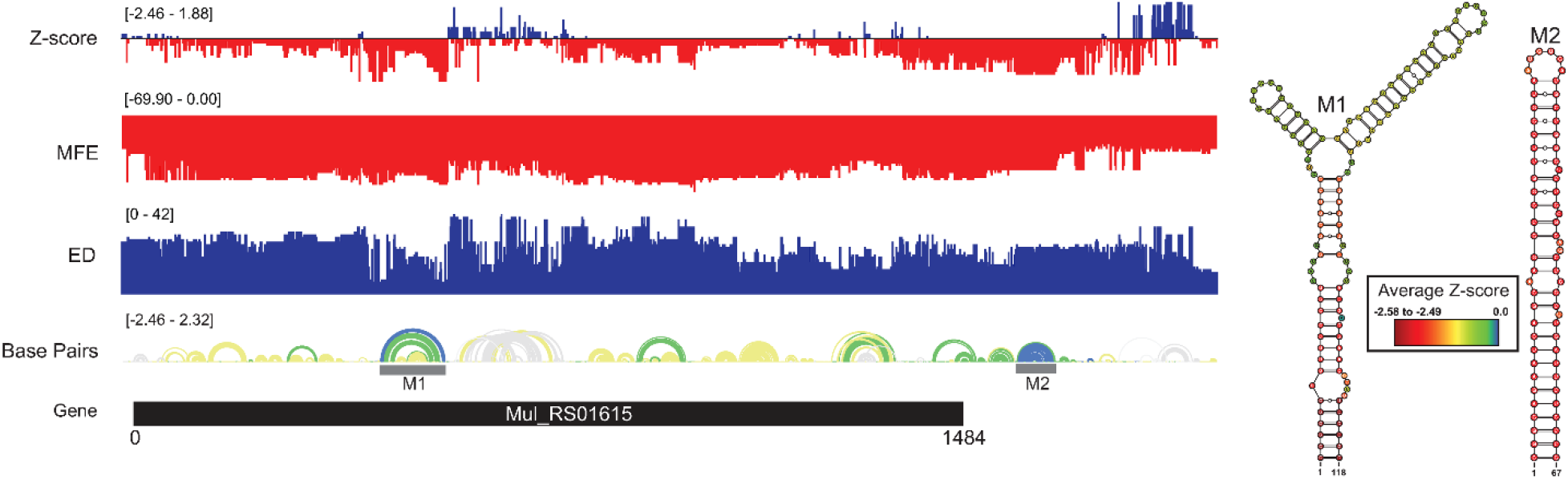
*Mul_RS01615* ScanFold results and 2D models. Global `ScanFold` results for *Mul_RS01615*. The ∆G z-score, minimum free energy (MFE), ensemble diversity (ED), gene cartoon, and base pair arc diagram (top to bottom) are shown to the left. The base pair arc diagram is annotated with gray boxes to show the location of M1-7 across the gene. All −2 ∆G z-score structures found are represented as 2D models to the right. The nucleotides of each structure are annotated with the average per nucleotide z-scores where the most negative are indicated in red and the most positive are indicated in blue.

### Analysis of antisense oligonucleotide accessibility

To explore the potential value of `ScanFold` data in identifying binding sites for short antisense oligonucleotides (ASO), we partitioned the `ScanFold-Scan` and `-Fold` results by averaging predicted metrics across short (18 nt) windows that approximate the size of potential ASO binding sites; this was further enhanced via predicting duplex binding affinities via the program `OligoWalk` (48) while considering the effects of `ScanFold` predicted local *intra*molecular structure on *inter*molecular duplex formation (all data available in **S3 File**). To facilitate analysis, these results were also plotted vs. `ScanFold-Fold` predicted base pairs (**S2 File**). Numerous short stretches of sequence across each gene of interest, including potentially accessible regions with positive z-scores, overlap strong predicted duplex binding sites. While the results *in toto* are potentially valuable for aiding in the identification and design of ASOs vs. *M. ulcerans*, we focus our attention on the two genes that have predicted ordered structures that encompass the start sites of translation, which are particularly attractive sites for ASO therapeutics.

The ASO accessibility results for the *desA2* homolog, *Mul_RS01365*, are summarized in **Fig. 7**. The trend toward enhanced thermodynamic stability of local RNA secondary structure in this gene is starkly illustrated, where significant dips in the partitioned averaged MFE and z-score overlap the translational stop site and `ScanFold-Fold` modeled low z-score structures. The inaccessibility of this region to ASOs is corroborated by `OligoWalk` predictions that show the overall duplex ΔGs are highly unfavorable: indeed, this region is predicted to have the most positive values across the gene. Conversely, the 5′ end, spanning the start codon, is predicted to have some of the least stable (least negative predicted average MFEs) local structure across the gene. Perhaps counterintuitively, there are dips in the average z-score and ensemble diversity that indicate ordered structure—indeed, `ScanFold-Fold` models show significantly ordered base pairing in this region (**Figs. 1** and **7**). Put another way, while the thermodynamic stability of this region is predicted to be low, it is still higher than expected given the nucleotide content, thus the ordered stability indicated by negative z-scores.

**Fig 7.**
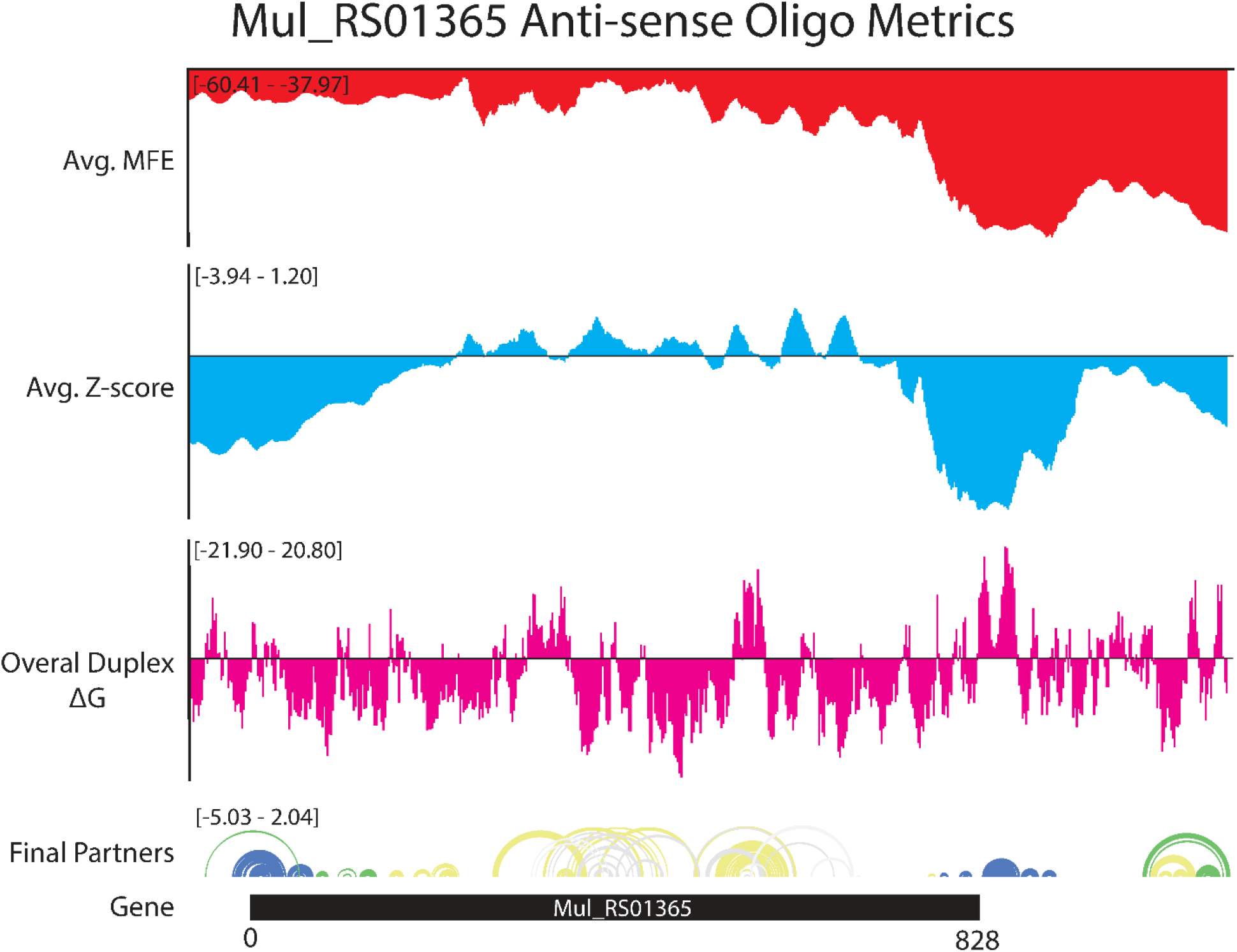
OligoWalk and ScanFold partitioned 18-mer data for *Mul_RS01365*. Data generated using `OligoWalk` and in-house script to partition `ScanFold` data into 18-mer averages for *Mul_RS01365*. Average MFE per 18-mer (red), average z-score per 18-mer (blue), overall duplex ΔG (pink), base pair diagram, and gene cartoon (top to bottom).

Favorable overall ASO duplex ΔGs span the start codon, despite it being contained in ordered structure (e.g., in M1, **Fig. 1**), indicating a potential for “strand invasion”, where an ASO can efficiently bind by breaking/replacing intramolecular helixes with intermolecular base pairs. To explore this further, and to illustrate how our data could facilitate ASO design, the most favorable (lowest ΔG, taking into account disruption of target structure) ASO predicted to occlude the start site was identified (**Fig. 8**). Here, an 18-mer ASO is predicted to bind to the mRNA with an overall ΔG of −9.8 kcal/mol, which takes into account the significant energy barrier (+15.8 kcal/mol) needed to disrupt 12 base pairs in the M1 hairpin structure. This disruption is predicted to totally ablate the terminal hairpin structure. Notably, other favorable ASO binding sites flank this optimal one (**Fig. 8** and **S3 File**) and thus, the ASO sequence can be extended in either direction to enhance predicted stability: e.g., to add additional stabilizing base pairs to the looped-out nucleotides predicted to be released upon disruption of the terminal hairpin loop.

**Fig 8.**
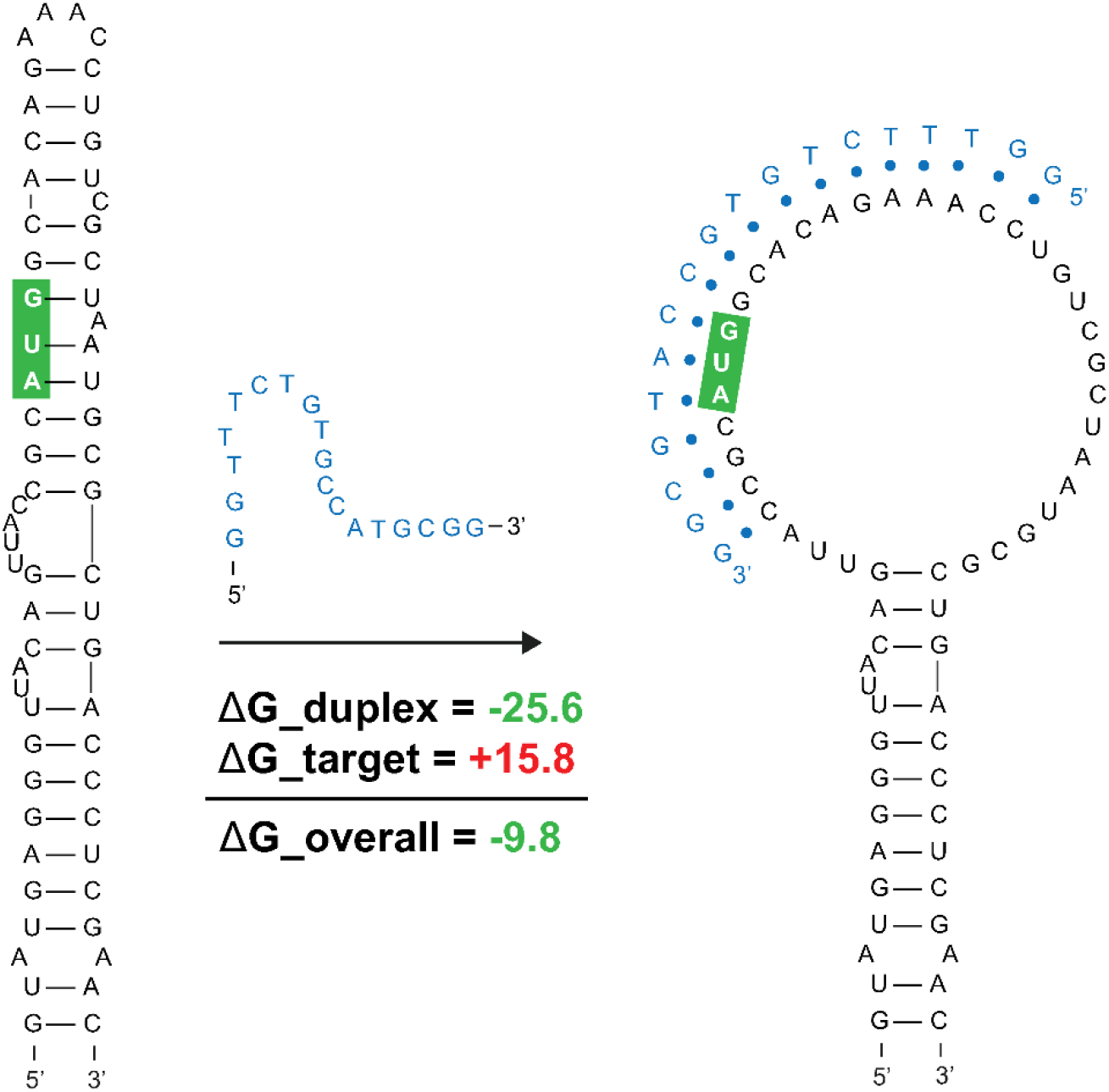
Predicted binding and strand invasion of *Mul_RS01365* Motif 1 by the most favorable 18-mer antisense oligonucleotide. The *Mul_RS01365* Motif 1 2D model, ASO of interest with `OligoWalk` data, and the predicted structure after strand invasion (left to right). Metrics in green indicate favorable binding, and the nucleotide outlined in green indicate the position of the start codon in the structure.

ASO accessibility results for the *rpoA* homolog *Mul_RS04730* are shown in **Fig. 9**. The partitioned MFE varies across the mRNA, however, it is predicted to be less stable in the 18-mers that span the start codon. Z-score is lowest toward the 5’ end of the RNA and steadily increases toward the 3’ end, indicating 18-mers are less likely to be engaged in ordered RNA structures as one moves along the sequence. Importantly, the overall predicted ASO duplex stabilities were most favorable in the region spanning the start codon: indeed, the most stable predicted duplex across the entire mRNA spans the start site (**Fig. 10)**. The overall predicted ASO duplex ΔG is −15.0 kcal/mol, which takes into account the relatively low barrier (+7.4 kcal/mol) needed to invade flanking hairpin structures in the target RNA (2 base pairs in each structure). Similar to *Mul_RS01365*, multiple other ASOs are predicted to bind near this optimal site, allowing for more stable sequences to be designed: e.g., by extending the optimal predicted ASO to further invade the small downstream hairpin structure.

**Fig 9.**
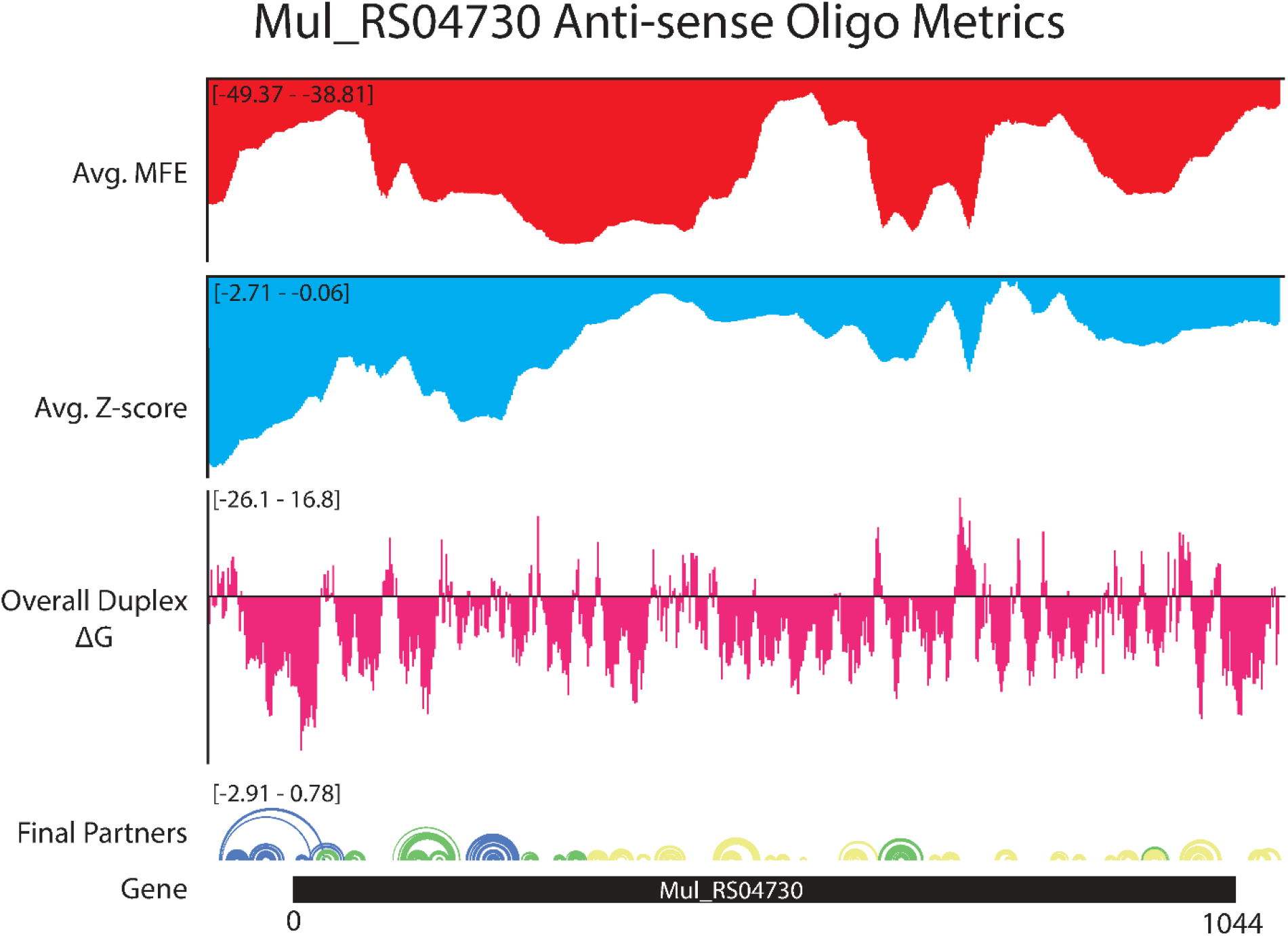
OligoWalk and ScanFold partitioned 18-mer data for *Mul_RS04730*. Data generated using `OligoWalk` and in-house script to partition `ScanFold` data into 18-mer averages for *Mul_RS04730*. Average MFE per 18-mer (red), average z-score per 18-mer (blue), overall duplex ΔG (pink), base pair diagram, and gene cartoon (top to bottom).

**Fig 10.**
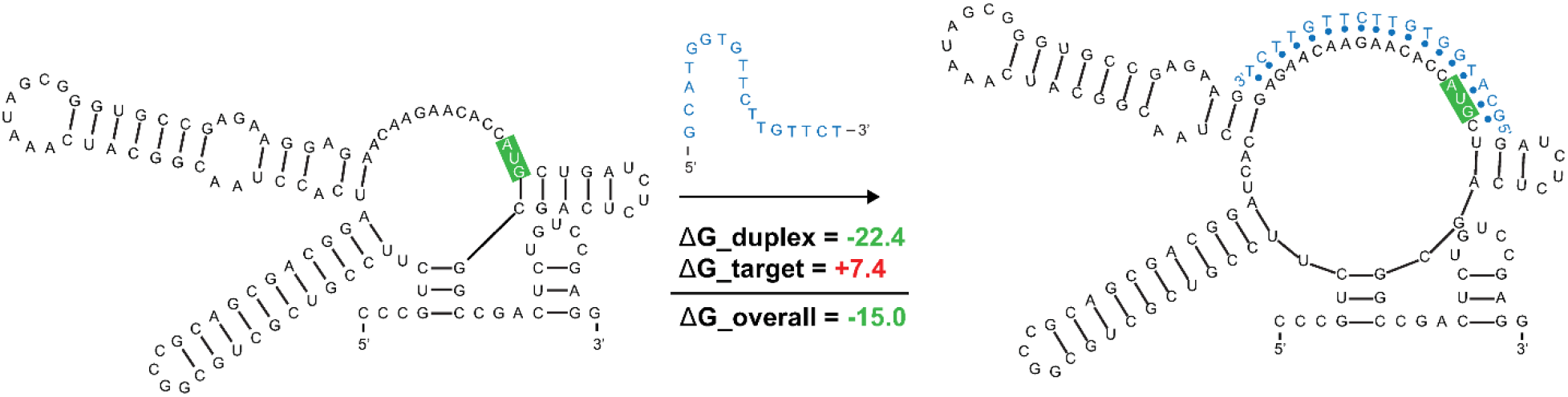
Predicted binding and strand invasion of *Mul_RS04730* Motif 1 by the most favorable 18-mer antisense oligonucleotide. The *Mul_RS04730* Motif 1 2D model, ASO of interest with `OligoWalk` data, and the predicted structure after strand invasion (left to right). Metrics in green text indicate favorable binding and the nucleotides outlined in green indicate the position of the start codon in the structure.

Beyond these interesting examples of ordered/structured RNAs that could be targeted with ASOs there are regions with apparent ordering for a lack of structure (positive z-scores) that could be especially accessible to ASOs (**S2 File**). For example, the 18-mer sequences that span the core coding region of *Mul_RS01365* are all predicted to have positive z-scores and relatively unfavorable MFEs (**Fig. 7**). This central portion of the coding region also possesses the most favorable predicted ASO binding sites for the gene, suggesting that this region is an especially attractive site for ASO binding.

## Discussion

This study reports the first comprehensive analysis of RNA secondary structure in *M. ulcerans*. Here, we focused on six genes based on their potential significance for drug targeting, and specifically adapted our structural analysis pipelines to obtain data that may provide valuable results for advancing *M. ulcerans* mRNAs therapeutics. Maps of the local RNA folding landscapes provide insights into the stability and potential for ordered/functional folding across each gene of interest. For example, looking at the `ScanFold` data for *Mul_RS01365* (both partitioned per-nucleotide and by 18-mers (**Figs. 1** and **7**, respectively)), we see thermodynamic stability (favorable/low MFE predictions) and ordered stability (low z-scores) at the 3′ end spanning the stop codon. This suggests potential roles for RNA structure and its thermodynamic stability in the termination of translation. Conversely, the coding region of this gene appears to be unstructured (as evidenced by mediocre MFEs and positive z-scores) perhaps to facilitate interactions with regulatory molecules or to promote rapid translation of this gene. Indeed, this latter idea is gaining traction as a mode for affecting protein folding (50): i.e., mRNA structural stability regulating the rate of translation to facilitate folding of proteins being translated.

While none of the analyzed genes had global biases for ordered structure (average z-score <−1; **Table 1**) clusters of ordered stability were present in each, where at least one defined structural motif could be predicted with exceptional (z-score < −2) base pairs, yielding 19 motifs in total across the six target genes. These sequences have been (presumably) ordered to fold over evolution and this proposition is supported by the overall conservation of secondary structure that was observed for each motif, as well as the identification of 4 statistically significant covarying base pairs. The functional roles of conserved, ordered RNA secondary structures in *M. ulcerans* and species with identified homologs can be diverse. For example, as noted above, structures in coding regions could affect translation rates. Structures may also be playing roles in modulating accessibility to regulatory molecules present in the bacterial cell or in mRNA turnover (e.g., modulating sensitivity to endogenous RNases). Significantly, two motifs span the translational start site: in *Mul_RS01365* and *Mul_RS04730* the start codons are modeled to lie within a helix and loop, respectively.

The instances where start sites are constrained within ordered structure present an interesting case for potential ASO therapeutics. Promising efforts at designing antimicrobial ASOs have focused on occluding the translational start sites of bacterial genes (51). For *Mul_RS01365* and *Mul_RS04730*, we predict oligos that can invade target intramolecular structure and, for *Mul_RS01365*, has the potential to greatly disrupt the ordered folding that may itself be functionally significant. That is to say, an ASO targeting the start site of *Mul_RS01365* may have two potential modes of action: obscuring the start site to impede translation and disrupting a secondary structure that may itself play roles in regulating translation. Beyond these two structural motifs, we present 17 others which may also be amenable to ASO-mediated disruption or that may be targeted with RNA-targeting small-molecules—a promising area of research for novel therapeutics. Notably, the 8 identified motifs found in *Mul_RS00210* are replicated multiple times in the virulence plasmid, which increases the concentration of potential ligand binding sites. Disrupting the genes responsible for mycolactone production could attenuate the pathogenicity of *M. ulcerans*. Our data also suggest regions of unusual thermodynamic *instability* across all analyzed genes, which may be ordered for accessibility to trans-regulatory molecules. These sites may be particularly attractive for ASOs that can mask such sites from interacting partners and/or stimulate mRNA degradation.

## Conclusion

To conclude, we present the first in-depth analysis of RNA structure in six key genes of *M. ulcerans*, the microbe responsible for the neglected tropical disease, Buruli Ulcer. Our results are made public to advance a basic understanding of the RNA biology of this pathogen—by providing conserved structural motifs of high likelihood of function (which may, themselves, serve as potential therapeutic targets). As well, we hope to advance novel therapeutics vs. *M. ulcerans* by providing data to facilitate antisense oligonucleotide design.

## Acknowledgements

This research was supported by National Institute of General Medical Sciences R01GM133810 to WNM and F31CA257090 to WBR. We would also like to thank the Science and Engineering Research Program (SERP) at Staten Island Technical High School led by Dr. John Davis.

## Supporting Information

**S1 File: M. ulcerans genomic data used in IGV-ScanFold.**

This file contains the M. ulcerans bacterial genome fasta, virulence plasmid fasta, and their associated gff3 genome annotations.

**S2 File: OligoWalk and 18-mer ScanFold bar charts**

This file contains the OligoWalk and 18-mer partitioned ScanFold data as bar charts overlaid against the gene cartoon for all six genes studied.

**S3 File: OligoWalk and 18-mer ScanFold raw data**

This file contains the raw output data from OligoWalk and in-house script for portioning 18-mer ScanFold data for all six genes of interest.

**S4 File: All ScanFold-Scan data**

This file contains a single folder for each gene studied. These folders contain the raw ScanFold-Scan output data such as per nucleotide MFE, ED, z-score, input, and output fasta files, and out file.

**S5 File: All ScanFold-Fold data**

This file contains a single folder for each gene studied. These folders contain the raw ScanFold-Fold output data such as the log file, base pair track, final partners data, all dot bracket files, all CT files, extracted structures gff3 file, and the global VARNA 2D model.

**S6 File: All cm-builder covariation data**

This file contains all the data required to run cm-builder and all the output files generated by INFERNAL and R-Scape.

**S1 Fig: All Mul_RS04200 ScanFold results**

Figure showing ScanFold results for Mul_RS04200 including z-score, MFE, ED, base pair diagram, gene cartoon, and 2D model of the structure with a z-score <−2.

**S2 Fig: All Mul_RS09540 ScanFold results**

Figure showing ScanFold results for Mul_RS09540 including z-score, MFE, ED, base pair diagram, gene cartoon, and 2D model of the structure with a z-score <−2.

